# Reciprocating RNA Polymerase batters through roadblocks

**DOI:** 10.1101/2023.01.04.522798

**Authors:** Jin Qian, Allison Cartee, Wenxuan Xu, Yan Yan, Bing Wang, Irina Artsimovitch, David Dunlap, Laura Finzi

**Author notes:** Contributing authors (Jin Qian, Allison Cartee, Wenxuan Xu, Yan Yan, Bing Wang, Irina Artsimovitch, David Dunlap).

## Abstract

RNA polymerases (RNAPs) must transit through protein roadblocks to produce full-length RNAs. Here we report real-time measurements of *Escherichia coli* (*E. coli*) RNAP passage through different barriers. As intuitively expected, assisting forces facilitated, and opposing forces hindered, RNAP passage through LacI bound to natural operator sites. Force-dependent differences were significant at magnitudes as low as 0.2 pN and were abolished in the presence of GreA, which rescues backtracked RNAP. In stark contrast, opposing forces promoted passage when the rate of backtracking was comparable to, or faster than the rate of dissociation of the roadblock, particularly in the presence of GreA. Our experiments and simulations indicate that RNAP may transit after roadblocks dissociate, or undergo cycles of backtracking, recovery, and ramming into roadblocks to pass through. We propose that such reciprocating motion also enables RNAP to break protein-DNA contacts holding RNAP back during promoter escape and RNA chain elongation, facilitating productive transcription *in vivo*.

## 1 Introduction

A dense array of proteins relevant to structure and function are associated with genomic DNA. These DNA-binding proteins play regulatory roles in cellular processes, such as genome packaging, and the recruitment and modulation of processive enzymes for replication and transcription [1–3]. These regulatory DNA-binding proteins vary significantly in their affinity for specific or non-specific DNA sequences, which may further change with physiological conditions [4]. Therefore, a motor enzyme, such as RNAP, encounters both low-and high-affinity DNA-bound roadblocking proteins. For uninterrupted transcription, these roadblocks must be transiently displaced during passage, whereas stalled RNAPs may require termination factors or convoys/collisions between polymerases to clear a template [5, 6].

Although roadblocks that interfere with transcription have been studied for decades, how RNAP surpasses a roadblock is known only for specific cases [7]. In principle, RNAP might passively wait for the roadblock to spontaneously dissociate, or it might actively dislodge the roadblock. Previous studies have shown that, when an elongation complex (EC) encounters an obstacle, such as a DNA lesion or a DNA-bound protein, it slides backwards [8]. In a backtracked EC, the nascent RNA occludes the active site, thereby blocking nucleotide addition and creating an obstacle. The arrested EC can be rescued by a trailing RNAP, a translating ribosome, a DNA translocase Mfd, which pushes RNAP forward, or by Gre factors, which facilitate the RNAP-mediated cleavage of the nascent RNA to restore the RNA 3’ OH in the active site [9–11]. The stability of the backtracked EC at a roadblock can also modulate the probability and/or rate of passage [12]. These findings were mostly based on run-off transcription assays that do not reveal the dynamics of roadblocked RNAP.

To reveal these dynamics, and expose general principles underlying RNAP progress through roadblocks, we used magnetic tweezers to monitor *E. coli* EC progress on DNA templates with either of two roadblock proteins, *lac* repressor (LacI) bound at sites with different affinities for the protein, or the mutant endonuclease EcoRI Q111, which binds but does not cut DNA. The experiments were conducted with up to 5 pN forces opposing or assisting RNAP translocation with or without GreA, the major anti-backtracking factor in *E. coli*. Our results indicate that RNAP can employ both passive and active mechanisms to overcome obstacles and that force and GreA modulate the pathway choice. We propose a model that well explains these observations and reveals the mechanism of ECs transiting through roadblocks.

## 2 Results

### 2.1 GreA and forces opposing or assisting RNAP translocation change pausing at roadblocks

Pauses at roadblocks during transcription elongation were measured using digoxigenin-end-labeled DNA templates containing a T7A1 promoter, a binding site for either the LacI protein: Os (*K_d_* = 10 fM), O1 (*K_d_* = 0.05 nM) or O2 (*K_d_* = 0.1 nM), or the EcoR1 Q111 protein (*K_d_* = 5 fM), and a *λ*T1 terminator (Figure 1A) [13–15]. Biotin-labeled RNAP holoenzyme coupled to a streptavidin-coated magnetic bead was introduced in flow chambers containing tethered DNA templates and manipulated in a magnetic tweezer microscope (Figure 1B). Whether force opposed or assisted transcription depended on which end of the template was digoxigenin-labeled and fixed to the glass, while the magnitude of the external force applied to RNAP was set by the separation between the permanent magnets above the flow cell and the sample.

**Fig. 1.**
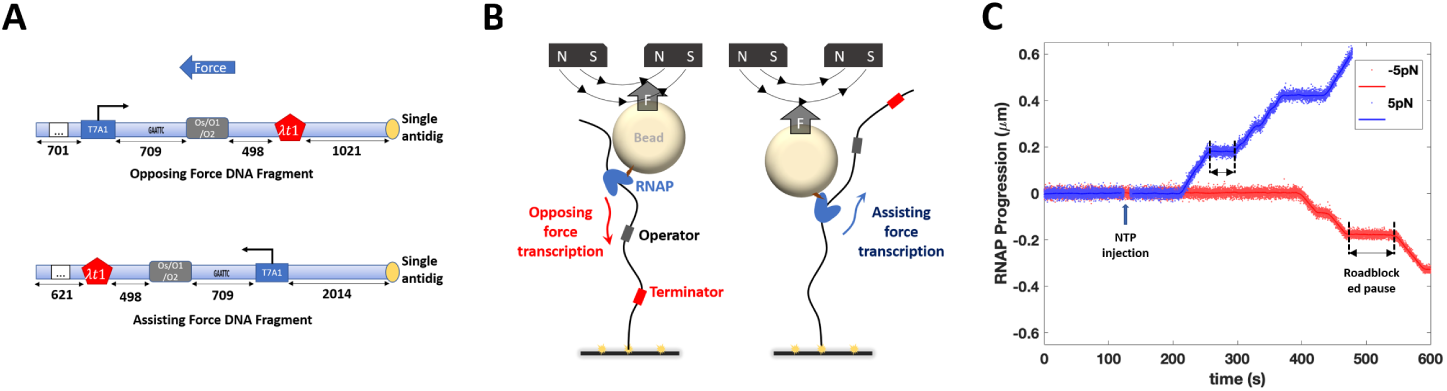
Features of the real-time experiments. (**A**) DNA templates for opposing and assisting force experiments have identical transcribed sequences. The numbers indicate the distances in base pairs. (**B**) A schematic illustration shows force opposing (left) or assisting (right) transcription. (**C**) Representative records of transcription template length as a function of time under opposing (red) and assisting (blue) force conditions show pauses at LacI roadblock sites (bracketed by black dashed lines).

Halted ECs were prepared as described in the Material and Methods section. RNA synthesis was restarted by the addition of all four NTPs (1 mM) at forces of 0.2, 0.7, 2.0, or 5.0 pN. These forces would generate from sub-kT to *∼*2kT of energy on a ratchet-like transcribing RNAP, well within the physiological range of tension experienced by the chromosomal DNA and the associated proteins [16]. In the presence of LacI, ECs paused near the operator site, where LacI was expected to bind, but eventually transited through the roadblock (Figure 1C). Different beads, due to variations in size and iron oxide content, exert slightly different forces producing different tether extensions. Thus, individual records of transcription (tether length *versus* time) were shifted and scaled to align them (Figure S1A and C). A step-wise fitting algorithm was then applied to produce a dwell-time histogram with distinct peaks at significant pauses and roadblocks (Figure S1B and D). The roadblock-associated pauses were significantly longer than ubiquitous, random pauses and were identified as occurring within ±20 nm (60 bp) of the roadblock binding site. A minor population (*<*10%) of traces had no roadblock-associated pauses, likely due to incomplete binding of roadblock proteins to the DNA tethers in the microchamber. Pauses shorter than 10 seconds were treated as ubiquitous pauses and were excluded from the analysis.

In the presence of LacI, we used DNA templates containing one of three *lac* repressor binding sites Os, O1 and O2, listed here in order of decreasing affinity (see above). For templates containing the O1 or O2 sites, nearly all ECs successfully transited through the roadblock within one hour. The pause times were distributed exponentially under all conditions except for LacI-O2 with assisting force, which will be discussed below, and the lifetimes changed with the direction (assisting *versus* opposing), but not the magnitude, of force (Figure 2A and B). As expected, the intermediate affinity LacI-O1 roadblocks produced longer pause times than the lowest affinity LacI-O2 roadblocks. On templates containing an artificial, symmetric binding site Os that binds LacI with the highest affinity, a fraction of ECs paused indefinitely at road-blocks (Figure 3A). Therefore, we determined both pause lifetimes (Figure 3B) and the fraction of RNAP passage through the LacI-Os roadblock (Figure 3C). Consistent with the LacI-O1/O2 induced pause times, LacI-Os induced pause times and passage ratio changed with the direction of force, whereas the magnitude of the force had a negligible effect.

**Fig. 2.**
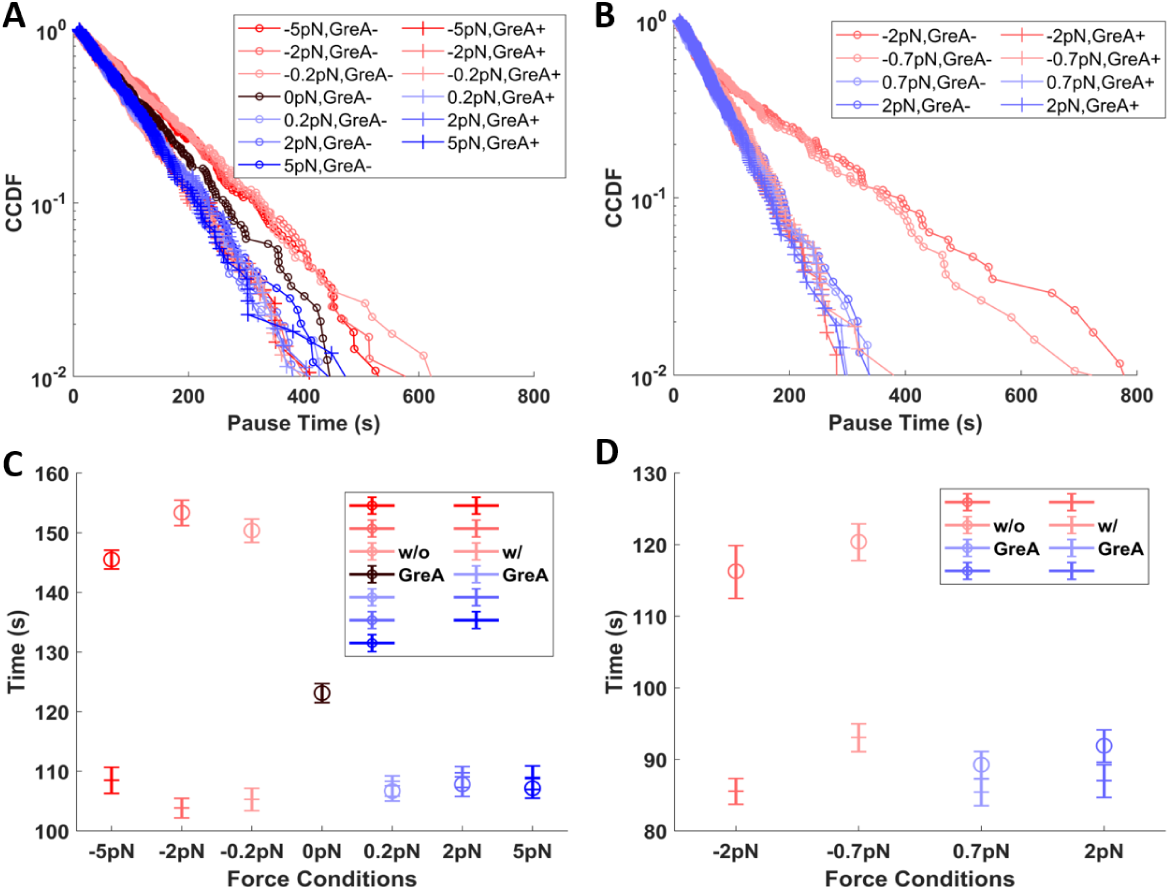
Pauses at LacI-O1 and LacI-O2 roadblocks under different force and GreA conditions. (**A**) The complementary cumulative distribution function (fraction of events longer than a given pause time, CCDF) and (**C**) characteristic times of pauses at LacI-O1 roadblocks shows that the longest pauses were associated with opposing force without GreA, followed by shorter pauses with no force, and even shorter pauses with assisting force or opposing force with GreA. (**B**) The CCDF and (**D**) characteristic times of pauses at LacI-O2 roadblocks shows that the longest pauses were associated with opposing force without GreA followed by shorter pauses with assisting force or opposing force with GreA.

**Fig. 3.**
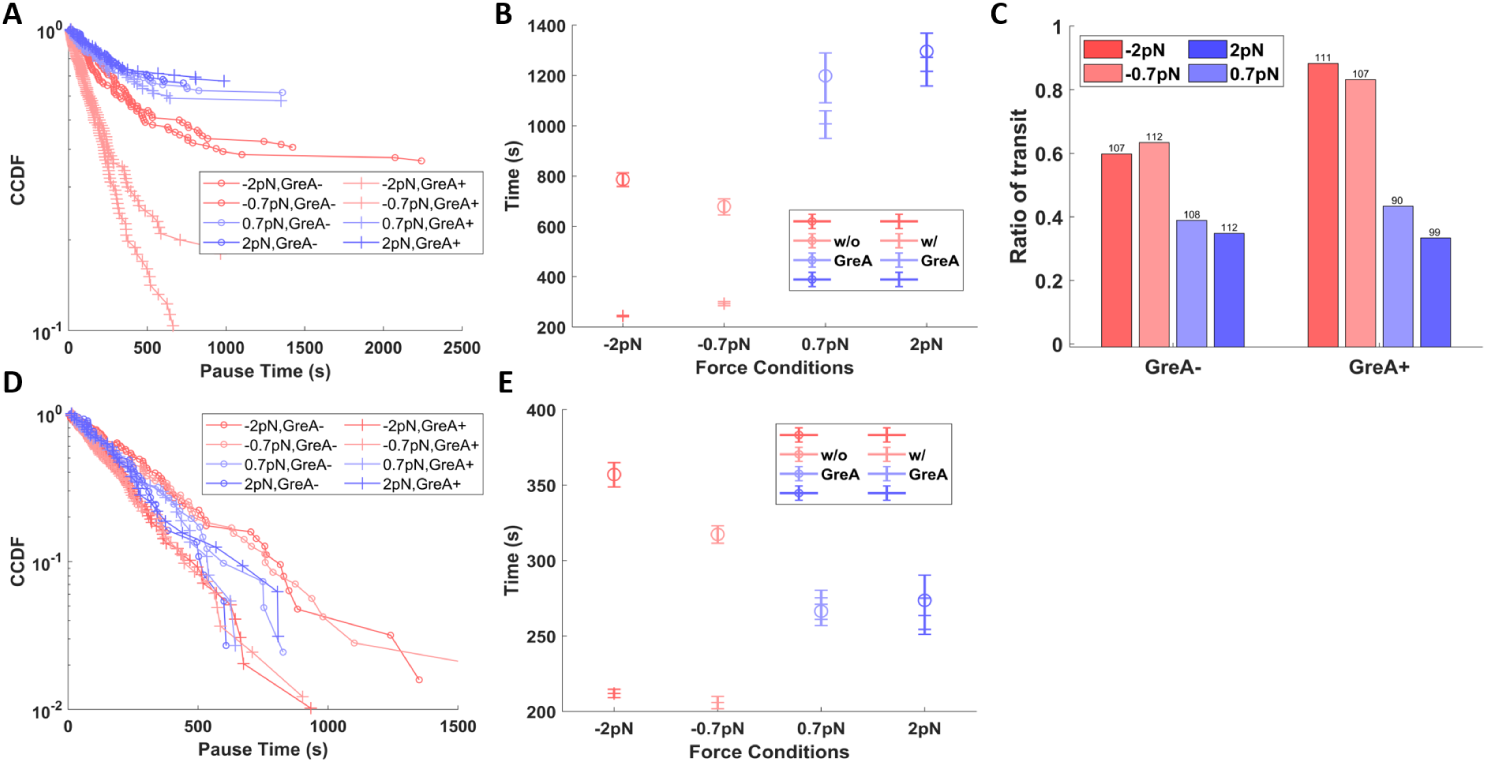
Pause times at and fraction of passage through LacI-Os roadblocks under different force and GreA conditions. (**A**) The CCDF and (**B**) characteristic times of all recorded pauses at LacI-Os roadblocks shows that the longest pauses were associated with assisting force with and without GreA, followed by shorter pauses with opposing force without GreA, and even shorter pauses with opposing force plus GreA. Note that all distributions except that for opposing force plus GreA include a significant fraction of indefinitely paused ECs. (**C**) Passage through LacI-Os roadblocks was more frequent under opposing than assisting force and was enhanced by the addition of GreA. (**D**) The CCDF and (**E**) characteristic times of pauses at LacI-Os roadblocks excluding those indefinitely shows that the longest pauses were associated with opposing force without GreA, followed by shorter pauses with assisting force without GreA, and even shorter pauses with opposing force plus GreA.

Backtracking by ECs has been widely associated with roadblocked transcription [9, 10]. Force that assists the EC translocation may affect pausing at roadblocks by preventing backtracking or by favoring recovery from backtracked states. Similarly, opposing force may promote backtracking or prevent recovery from backtracked states. To reveal the contribution of backtracking to pausing at roadblocks, GreA was added to rescue backtracked ECs by promoting the cleavage of the 3’ end of nascent RNA threaded through the active site [17]. GreA did not change pauses under assisting force (Figure 2C and D, 3B), most likely because EC backtracking, and therefore GreA activity, were negligible under assisting-force conditions. On the contrary, GreA apparently increased the rate of recovery from backtracked states to accelerate passage through roadblocks under opposing force (Figure 2C and D, 3B). This is consistent with previous work showing that GreA enhanced passage through the LacI roadblocks [18]. Remarkably, unlike sequence-induced backtracking [19], even a very gentle force of 0.2 pN significantly reduced backtracking by roadblocked ECs. This is consistent with the low energy barriers to backtracking found previously for Rpo41 and PolII [20].

### 2.2 GreA and tension reveal two paths through roadblocks

The direction of force and the addition of GreA changed pausing at LacI bound to O1, O2, or Os operators differently (Figures 2C and D, and 3B). For DNA templates containing either O1 or O2, opposing force lengthened pauses relative to the assisting-force baseline, but adding GreA restored that baseline. These results support an accepted notion that backtracking, without recovery promoted by GreA, inhibits productive RNA synthesis. Strikingly, however, on templates containing Os, opposing force without GreA shortened the pauses to a level below that observed under assisting force, and the effect was enhanced by the addition of GreA. The unexpected stimulatory effect of opposing force suggested that relatively fast cycles of backtracking and subsequent recovery, a reciprocating motion, may enable RNAP to batter through relatively slowly dissociating roadblocks, with GreA accelerating repeated collisions between RNAP and the roadblock. This could be particularly important if interactions with RNAP were to delay dissociation of the roadblock. Such an interaction was not anticipated, but lac repressor has been reported to bind to RNAP [21], and this interaction was confirmed by co-immobilization (Figure S2).

To test this hypothesis, transcription assays were performed against another strong roadblock, the mutant EcoRI Q111 endonuclease which does not interact with RNAP. The protein binds with high affinity but does not cut at 5’-GAATTC sites. As expected, EcoRI Q111 successfully blocked nearly 100% of ECs in standard buffer conditions independently of the force. The addition of GreA increased the level of passage through this roadblock to 20% only under opposing force (Figure 4A, 50 mM [KGlu]). Notably, adding GreA to a roadblocked EC produced passage shortly thereafter (Figure 4B), suggesting that opposing force and GreA likely promote transit through these long-lived roadblocks.

**Fig. 4.**
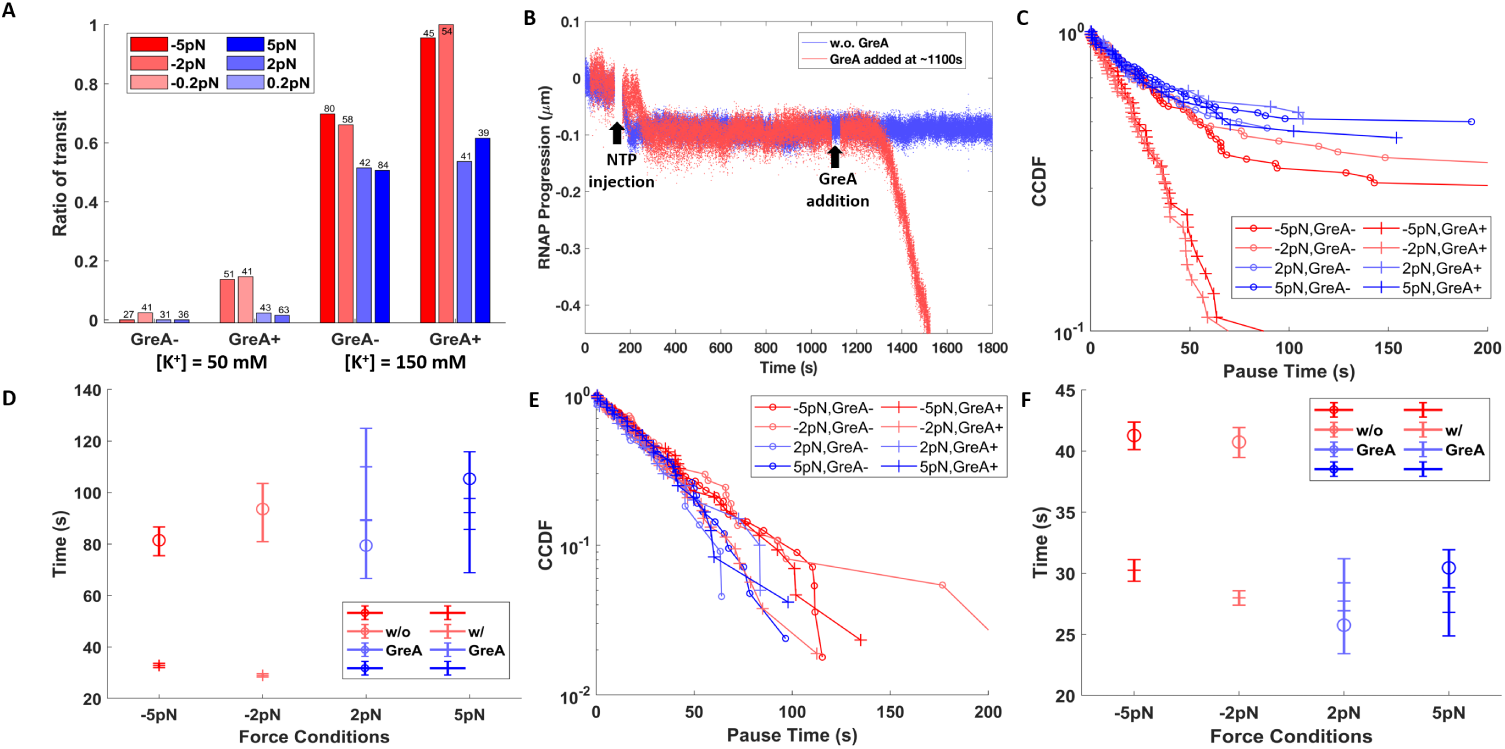
Pause times and the fraction of transit through EcoRI Q111 roadblocks under different force and GreA conditions. (**A**) Passage through EcoRI Q111 roadblocks was rare in 50 mM [KGlu] but increased dramatically in 150 mM [KGlu] especially upon the addition of GreA. (**B**) Without GreA most ECs in 50 mM [KGlu] buffer paused indefinitely at EcoRI roadblocks (blue), but adding GreA (red, *∼*1100 s) rescued paused ECs that resumed transcription (red, *∼*1300 s)). (**C**) The CCDF and (**D**) characteristic times of all recorded pauses at EcoRI Q111 roadblocks show that the longest pauses were associated with assisting force with and without GreA or opposing force without GreA, and shorter pauses with opposing force plus GreA. Note that all distributions except that for opposing force plus GreA include a significant fraction of indefinitely paused ECs. (**E**) The CCDF and (**F**) characteristic times of pauses including only ECs that eventually pass through EcoRI Q111 roadblocks in 150 mM [KGlu].

Under increased salt concentration (150 mM [KGlu]), the affinity of EcoRI Q111 protein for the recognition site weakens such that a considerable fraction of ECs passed through roadblocks (Figure 4A, 150 mM [KGlu]) [22]. Therefore, we measured both the distribution of roadblock-induced pause times and the fraction of passage by ECs. We note that a considerable fraction of ECs dissociated before transiting through EcoRI Q111 roadblocks under high salt (data not shown) and were categorized as indefinitely blocked ECs, even though the mean assisting-force pause lifetime was only *∼*80 seconds, which is shorter than pauses at LacI-O1 and comparable to pauses at LacI-O2 (Figure 4C and D). We postulate that the elevated salinity induced dissociation of ECs at roadblocks, preventing observation of more passage events. The addition of GreA reduced the opposing force pauses to a level below the assisting-force baseline level (Figure 4D). Opposing force also increased the fraction of ECs transiting through EcoRI Q111 roadblocks, which increased even further with the addition of GreA (Figure 4A, 150 mM [KGlu]). The observed pause lifetime agrees with the estimation of the dissociation constant of EcoRI Q111 under high salinity *K_d_* = 0. 12 nM, assuming a linear relationship between *ln*(*K_a_*) and *ln*([*M* ^+^]) [23].

A comparison between LacI-O2/O1 induced pauses and LacI-Os/EcoRI Q111 induced pauses shows two clearly different responses of ECs to changes in the direction of force and the addition of GreA. For the LacI-O2/O1 roadblocks, assisting force, either with or without GreA, sets a baseline pause before transit, and GreA must be added to reach that baseline under opposing force. On the contrary, for LacI-Os and EcoRI Q111 roadblocks, opposing force may hasten passage more than the assisting force, especially when GreA is present. To test if the different responses are associated with roadblock strengths, subsets of LacI-Os and EcoRI Q111 data, which exclude the indefinitely stalled ECs, were analyzed. This selects a sub-population of ECs paused at shorter lived, relatively lower affinity roadblocks. For such LacI-Os roadblocks, opposing force produced pauses longer than the assisting-force baseline, while addition of GreA shortened the opposing pauses significantly to a level below that baseline (Figure 3D and E). Similarly, for EcoRI Q111 roadblocks of short duration, the addition of GreA shortened the opposing-force pause time to the level of the assisting-force baseline (Figure 4E and F), as observed for LacI-O1 and LacI-O2 roadblocks. These results confirm that the lifetime of the roadblock can bias the transit efficiency under opposing forces relative to the assisting force baseline.

The fact that pauses at roadblocks under assisting force are insensitive to GreA indicates that assisting force prevents backtracking. ECs must therefore follow a passive pathway, remaining transcriptionally active, ready to proceed when roadblocks dissociate. For LacI-O1 and LacI-O2, this passive pathway appears to be an efficient transit mechanism; while for LacI-Os and EcoRI Q111 roadblocks, this pathway leads to a significant portion of indefinitely stalled ECs (Figure 3C, 4A).

In contrast, GreA significantly enhances transit through roadblocks under opposing force. This finding is consistent with an active, reciprocating pathway that involves backtracking of ECs and GreA-enhanced recovery. Notably, the analyses of both pause lifetimes and fractions of transit suggests that the active pathway speeds transit through LacI-Os and EcoRI Q111 roadblocks (Figure 3B and C, 4C and D) while slowing transit through LacI-O1 and LacI-O2 roadblocks (Figure 2C and D). The addition of GreA shortened the opposing force pause times at different roadblocks by different amounts, suggesting that the active pathway may involve multiple cycles of backtracking and recovery before ECs successfully batter through roadblocks.

### 2.3 The effects of force and GreA fit a hybrid transit model

A hybrid model including the reciprocating and passive pathways is consistent with the data. Figure 5A depicts the progression through roadblocks via different states along these pathways. The passive pathway progresses through states and the active pathway through ○1 *→*(○2 *→*○3 *→*○2) *_n_→*○5 *→*○6 with *n* cycles of back-tracking and recovery. The model includes three kinetic parameters *k*_1_, *k*_2_ and *k*_3_, which represent the backtrack rate, backtrack recovery rate, and roadblock dissociation rate, respectively. Parameter *P* 1 represents the probability of dislodging the roadblock at each encounter. Therefore, the transit rate of the passive pathway is simply *k*_passive_ = *k*_3_, and rate of active pathway is *k*_active_ = *P*_1_ *k*_1_*/*(1 + *k*_1_*/k*_2_).

**Fig. 5.**
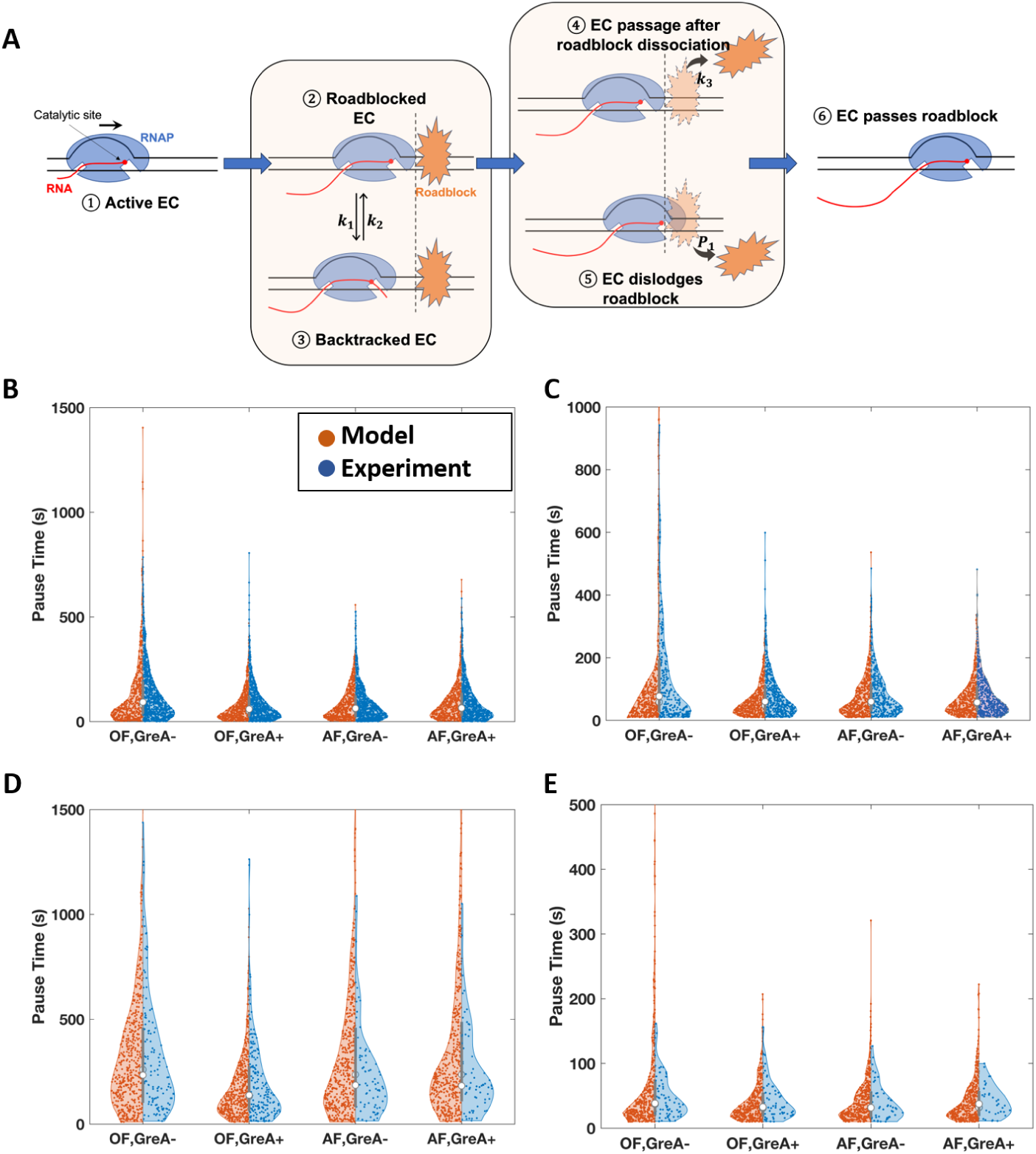
Model and simulation results. (**A**) A proposed model of EC transit through a roadblock includes six states: State ①: Transcription prior to the encounter of roadblock; ②: An EC encounters the roadblock; ③: At a roadblock an EC backtracks with a backtracking rate *k*_1_ and recovery rate *k*_2_; ④: A roadblock dissociates from DNA spontaneously with a dissociation rate *k*_3_; ⑤: An actively transcribing EC, including a recently backtracked EC, has a probability *P*_1_ of dislodging the roadblock; ⑥: An EC transits through the roadblock either by actively dislodging the roadblock or after spontaneous dissociation of the roadblock. (**B**) - (**E**) Simulations produced pause time distributions very similar to those observed in LacI-O1, LacI-O2, LacI-Os and EcoRI Q111 experiments under different conditions. ***OF***, opposing force, ***AF***, assisting force.

Notice that ECs likely execute multiple cycles of backtracking and recovery before successfully dislodging the roadblock, which may also spontaneously dissociate during these cycles. Since an EC moves stochastically through the various states of the two pathways to pass through a roadblock, a Monte Carlo simulation is suitable to reproduce distributions of pause times at roadblocks in different conditions. Indeed, such a simulation faithfully reproduced the effects of opposing *versus* assisting force and the addition of GreA (Figure 5B-E). Details of the simulation are described in Materials and Methods.

The model predicts changes in pause times and transit percentages produced by GreA and changes in the direction of force. The effects of force and GreA are determined by the relative kinetics of RNAP and roadblock proteins [24], specifically the rate at which cycles of backtracking-recovery occur (*k*_active_) *versus* the rate of roadblock dissociation (*k*_passive_). The simulation reveals three distinct regimes. If cycles of backtracking-recovery are much slower than the dissociation of roadblocks, (passive route regime: *k*_active_ *≪ k*_passive_), an EC nudged into a backtracking-recovery cycle by opposing force, may not finish a cycle before spontaneous dissociation of the road-block. For such conditions, the model simulated identical relatively short pauses for the assisting/Gre-, assisting/Gre+ and opposing/Gre+ conditions, but relatively long pauses for the opposing/Gre- condition, resembling the results from the LacI-O2, LacIO1 and the low-affinity sub-population of high salinity EcoRI Q111 measurements (Figure S3A, Figure 2C and D, 4F). Notably, in this regime, the simulated pause distribution for the opposing/Gre- condition is much better fitted by a double exponential, representing two stochastic processes with distinct rate constants. A similarly long tail in the distribution of pauses at LacI-O2 roadblocks in opposing/Gre- conditions but not in other experimental conditions, further supports the model (Figure 2B).

When the two rates are comparable, (hybrid route regime: *k*_active_ *∼ k*_passive_), opposing force could either lengthen dwell times if RNAP is backtracked at the moment of roadblock dissociation, or shorten dwell times if RNAP successfully dislodges a road-block before it spontaneously dissociates. The simulated pause distribution (Figure S3B) resembles the distribution observed with EcoRI Q111 roadblocks at high salt and the sub-population of low-affinity LacI-Os roadblocks, for which opposing/Gre- led to longer pauses than in assisting force conditions and opposing/Gre+ led to shorter pauses (Figure 3E, 4D). In this regime, the active and passive pathways proceed at similar rates, and all distributions are fitted well by a single exponential.

Finally, if roadblock dissociation rate is relatively slow compared to backtracking and recovery cycling, (active route regime: *k*_active_ *≫ k*_passive_), the passive pathway is inefficient, the active pathway is more successful, and opposing force favors backtracking-recovery cycles to produce shorter pauses and higher fractions of EC transit, especially with GreA present. The simulated results (Figure S3C) concur with the experimental results for LacI-Os and low salt EcoRI roadblocks for which opposing force produced shorter pauses and higher fractions of transit than assisting force, and GreA further shortened the pauses and further increased the fraction of transit (Figure 3B and C, Figure 4A).

Table 1 summarized the values of kinetic parameters by fitting the model to the experimental results. As expected, the passive route rate agrees with the affinity of these roadblocks, *k*_passive_(high salt EcoRI) *∼ k*_passive_(O2) *> k*_passive_(O1) *> k*_passive_(Os). Moreover, the model suggests that the transition through EcoRI Q111 (high salt) and LacI-Os roadblocks occur in the hybrid route regime, *k*_active_(high salt EcoR1/Os) *∼ k*_passive_(high salt EcoR1/Os), while transition through LacI-O1 and LacI-O2 roadblocks occurs in the passive route regime, *k*_active_(O1/O2) *≪ k*_passive_(O1/O2). These results indicate that cycles of backtracking and recovery occur faster when ECs confront EcoRI roadblocks than LacI roadblocks. We postulate that the different DNA sequences upstream of EcoRI and LacI binding sites may contribute to the difference in the rate of backtracking cycle. Indeed, the analyses of EC energy profiles indicate that EC is less stable upstream of the EcoRI roadblock, which might suggest faster backtracking-recovery cycles (Figure S4). Furthermore, the rate of the backtracking-recovery at a LacI roadblock may be slowed by the interaction between RNAP and LacI (Figure S2), so that battering through only becomes faster than the stable passive dissociation in the case of the LacI-Os roadblock.

**Table 1.** Kinetic parameters generated by fitting model to the experimental pause time distributions

| Parameters | LacI-O2 | LacI-O1 | LacI-Os | EcoRI (150 mM [KGlu]) |
| --- | --- | --- | --- | --- |
| $k_{\text{passive}} (s^{-1})$ | 0.016 | 0.009 | 8e-4 | 0.01 |
| $k_1 (s^{-1})$ | 0.01 | 0.01 | 0.01 | 0.04 |
| $k_2 (s^{-1})$ | 0.005 | 0.005 | 0.005 | 0.08 |
| $P_1$ | 0.2 | 0.2 | 0.2 | 0.2 |
| $k_{\text{active}} (s^{-1})$ | 6.6e-4 | 6.6e-4 | 6.6e-4 | 0.005 |

The experimental data and the model suggest that the characteristic time of a backtracking-recovery cycle is about 150 seconds for LacI roadblocks and about 40 seconds for EcoRI Q111 roadblocks in high salt, which is longer than the typical lifetime of sequence-induced, backtracked pauses [18, 19]. However, similar pauses are frequently observed in a variety of experiments and are classified as stabilized-backtracked pauses [25, 26]. Such force-independent, lengthy pauses support the hypothesis that roadblock-induced backtracking might include a force-independent, rate-limiting intermediate state.

## 3 Discussion

Our experimental data and simulations support a model in which RNAP can utilize two mechanisms to bypass obstacles on its path, dynamically shifting between alternative modes of transit through roadblocks of differing strength. In the passive mode, ECs wait and readily proceed upon roadblock dissociation. In the reciprocating mode, ECs execute cycles of backtracking and recovery to batter through roadblocks. Roadblock duration determines which pathway produces more passage. ECs primarily followed a passive route through LacI-O1 and LacI-O2 roadblocks, while the reciprocating backtracking-recovery pathway shortened pauses producing more frequent passage through LacI-Os and EcoRI Q111 roadblocks. Rather than being a mere nuisance, backtracking significantly helps EC evict DNA-bound proteins that dissociate relatively slowly making the passive route unproductive.

Unexpectedly, roadblocked ECs are exquisitely sensitive to mechanical force and as little as 0.2 pN can significantly change transit. Forces of such magnitude that could impact transcription are prevalent in physiological conditions due to genome anchoring and contacts [27, 28]. Therefore, the passage of ECs through roadblocks could be significantly biased by genome architecture and dynamics. However, the passage of ECs through roadblocks was also surprisingly insensitive to the magnitude of forces ranging from 0.2 pN to 5pN. This contrasts with previous reports in which greater opposing force led to longer backtracked pauses [29, 30]. We hypothesize that roadblock-associated backtracking involves a force-sensitive intermediate state followed by a force-insensitive rate-limiting step. Previous studies suggested that external forces bias the efficiency of backtracked pauses by modulating the forward transcription rate from backtracked positions, whereas bi-directional fluctuations of backtracked ECs were relatively resistant to external forces [19, 31]. We speculate that the force-insensitive rate-limiting step is likely the bi-directional diffusion of backtracked ECs, whereas the force-sensitive intermediate might be a state unique to ECs contacting roadblock proteins. The difference between roadblock-and sequence-induced back-tracks indicates that there might be functional and mechanistic differences. The former serves to dislodge roadblocks and is easily triggered, or prevented, by the change in external tension, while the latter functions as a universal error-correction mechanism and is more resistant to change. Meanwhile, the characteristic time of the roadblock-induced backtracks and recovery cycle is predicted to be 40–150 seconds, which is significantly longer than the lifetime of the sequence-induced backtracks and resembles the dwell time of the “stabilized-backtracked pauses” described elsewhere [26]. Elucidating the mechanisms and functions of backtracked pauses of different lifetimes remains an important subject for future biochemical studies.

Previous studies suggested that backtracked RNAPs contribute to transcription traffic jams. Indeed, *E. coli* employs multiple mechanisms to rescue backtracked RNAPs, such as Gre-factor-dependent RNA cleavage, transcription-translation coupling, and/or trailing RNAPs [6, 8, 32]. Our study reveals that backtracking may actually have a positive impact, in that an active, reciprocating pathway with backtracking-recovery cycles can promote efficient passage through relatively long-lived roadblocks, which is critical to prevent RNAP from being stalled and targeted by exonucleolytic activity [33]. Our study also explains the different behavior of

RNAP and helicase RecBCD when encountering the EcoRI roadblock. The latter can push the EcoRI roadblock thousands of base pairs before finally evicting it, whereas RNAP remains at the roadblock position until passage [34]. In contrast to ATP-driven helicases and translocases, RNAP likely exerts little chemo-mechanical force on nucleoprotein complexes due to the Brownian ratchet nature of translocation. Indeed, RNAP can only generate chemo-mechanical force during the incorporation of an NTP [35]. Therefore, repetitive backtracking-recovery cycles are most likely required to perturb and eventually displace roadblocks.

We note that hindrance by stable protein-DNA contacts is not limited to extrinsic proteins bound to DNA in the the path of RNAP. In fact, RNAP faces the same problem every time it initiates transcription: contacts between the promoter DNA elements and the initiation *σ* factor hold RNAP back, triggering repeated synthesis and release of short abortive RNAs or formation of arrested complexes [36]. Gre factors facilitate promoter escape [37], suggesting that cycles of backtracking and RNA cleavage are required to rupture *σ*-DNA interactions. Similarly, transcription elongation factors that are recruited to RNAP via DNA, such as RfaH, promote backtracking and depend on Gre factors for escape from the recruitment site [38]. Thus, cycles of backtracking, cleavage, and re-extension are likely to be required for uninterrupted RNA synthesis in all cases when strong DNA-protein contacts, either ahead or behind the moving RNAP, hinder RNA chain extension.

In summary, this study reveals a hybrid mechanism of passage through protein roadblocks along the path of transcription elongation complexes. ECs can passively wait for dissociation of roadblocks or actively batter through them, and backtracking may lengthen or shorten pauses at roadblocks depending on the longevity of the roadblocks and their interactions with ECs. The effects of tension and the transcript cleavage factor GreA demonstrate that structural (affinity) or dynamic (tension) factors can modulate the efficiency of these pathways and the route followed by ECs to pass through roadblocks. Various elongation factors *in vivo* might finely tune the efficiency of either pathway making them quite distinct to produce a deterministic response that serves specific biological purposes.

## 4 Materials and Methods

### 4.1 Preparation of Proteins

#### 4.1.1 *E. coli* Biotinylated RNAP Holoenzyme

*E. coli* BL21 (*λ*DE3) harboring pIA1202 (*α*-*β*-*β*’[AVI][His]-*ω*) was cultured in LB at 37*°*C. The expression was induced at OD600 *∼*0.5 with 0.5 mM IPTG for 5 hours at 30*°*C. To achieve better biotinylation, biotin ligase (BirA; Addgene#109424) was expressed in *E. coli* BL21 (*λ*DE3) in the same way. Cells were pelleted by centrifugation (6000 x g, 4*°*C, 10 min). Cell pellets were mixed and resuspended in Lysis Buffer (10 mM Tris-OAc pH 7.8, 0.1 M NaCl, 10 mM ATP, 10 mM MgOAc, 100 *µ*M d-biotin, 5 mM *β*-ME) supplied with Complete EDTA-free Protease Inhibitors (Roche) per manufacturer’s instructions.

Cells were sonicated and cell debris was pelleted by centrifugation (20,000 x g, 40 min, 4*°*C). The cleared cell extract was incubated with Ni Sepharose 6 Fast Flow resin (Cytiva) for 40 min at 4*°*C with agitation. The resin was washed with Ni-A Buffer (25 mM Tris pH 6.9, 5% glycerol, 150 mM NaCl, 5 mM *β*-ME, 0.1 mM phenylmethylsulfonyl fluoride (PMSF) supplemented with 10 mM, 20 mM, and 30 mM imidazole. Protein was eluted in Ni-B Buffer (25 mM Tris pH 6.9, 5% glycerol, 5 mM *β*-ME, 0.1 mM PMSF, 100 mM NaCl, 300 mM imidazole).

The sample was diluted 1.5 times with Hep-A Buffer (25 mM Tris pH 6.9, 5% glycerol, 5 mM *β*-ME) and then loaded onto Heparin HP column (Cytiva). A linear gradient between Hep-A and Hep-B Buffer (25 mM Tris pH 6.9, 5% glycerol, 5 mM *β*-ME, 1 M NaCl) was applied. The biotinylated RNAP core is eluted at *∼*40 mS/cm.

The elution from Heparin HP column was diluted 2.5 times with Hep-A Buffer and loaded onto Resource Q column (Cytiva). A linear gradient was applied from 5% - 100% Hep-B Buffer. The biotinylated RNAP core was eluted at *∼*25 mS/cm.

Fractions from the elution peaks were analyzed by SDS-PAGE. Those containing purified protein were combined and dialyzed against Storage Buffer (20 mM Tris-HCl, pH 7.5, 150 mM NaCl, 45% glycerol, 5 mM *β*-ME, 0.2 mM EDTA).

*E. coli* RNAP holoenzymes were reconstituted from *σ*^70^ initiation factor, expressed as previously described [39], and the biotinylated RNAP core enzyme.

#### 4.1.2 Lac Repressor Protein

Lac repressor protein (LacI) was provided by Kathleen Matthews (Rice University).

#### 4.1.3 EcoRI Q111 Protein

*E. coli* BL21 (*λ*DE3) harboring pVS9 (His6-tagged EcoRI Q111) was cultured in LB at 37*°*C. The expression was induced at OD600 *∼*0.8 with 0.3 mM IPTG for 3 hours at 37*°*C. Cultures were chilled on ice for 15 min and cells were pelleted by centrifugation (6000 x g, 4 oC, 10 min).

The cell pellet was resuspended in Lysis Buffer (50 mM Tris-HCl, pH 7.5, 1.5 M NaCl, 5% glycerol, 1 mM *β*-ME) supplemented with 1 mg/ml lysozyme, 0.1% Tween 20, and Complete EDTA-free Protease Inhibitors (Roche) per manufacturer’s instructions. The suspension was incubated on ice for 45 min and sonicated to disrupt cells. Cell debris was pelleted by centrifugation (20,000 x g, 30 min, 4*°*C). Cleared extract was incubated with Ni-NTA Agarose slurry (Qiagen) for 30 min at 4*°*C with agitation. After washing with Wash Buffer (50 mM Tris-HCl, pH 7.5, 0.5 M NaCl, 5% glycerol, 1 mM *β*-ME), elution was carried out with 20 mM, 50 mM, 100 mM, and 300 mM imidazole in Heparin Buffer (50 mM Tris-HCl, pH 7.5, 200 mM NaCl, 5% glycerol, 1 mM *β*-ME). Fractions containing EcoRI Q111 were pooled and loaded onto Heparin HP column (Cytiva). Protein was eluted by a linear gradient from 200 mM to 800 mM NaCl in Heparin Buffer.

Fractions from the elution peaks were analyzed by SDS-PAGE. Those containing purified protein were combined and dialyzed against Storage Buffer (20 mM Tris-HCl, pH 7.5, 300 mM KCl, 50% glycerol, 0.2 mM DTT, 1 mM EDTA).

#### 4.1.4 Gre Factor

Gre factors were purified from plasmids constructed in the Artsimovitch lab as previously described [40], analyzed by SDS gel electrophoresis, and tested for RNA cleavage activity. These purified proteins have been crystallized and used in functional and single molecule studies by many groups.

### 4.2 Transcription Templates for Magnetic Tweezers Assays

All DNA fragments for MT experiments were PCR amplicons from plasmid templates pDM E1 400, pDM N1 400, pZV NI 400 or pDM N2 400, single-digoxigenin labeled forward and unlabeled reverse primer pairs, and Q5 Hot Start High-Fidelity 2X PCR Master Mix (New England Biolabs). The transcribed region had the following spacings: Promoter-709 bp-Lac operator-498 bp-Terminator, with an EcoRI binding sequence GAATTC between the promoter and the operator site, as illustrated in Figure 1A. For the opposing force experiments, primers 5’-dig-ATCGTTGGGAACCGGAG and 5’- AGCTTGTCTGTAAGCGGATG were used to generate 3k bp DNA fragments with 1021 bp between the chamber surface anchor point and the transcription start site. For the assisting force experiments, primers 5’-dig-GCTTGGTTATGCCGGTACTG and 5’-ACGACCTACACCGAACTGAG were used to generate 4k bp DNA fragments with 2014 bp between the anchor point and transcription start site. The longer separation in the assisting force DNA fragment reduces adhesion of DNA-attached magnetic beads to the chamber surface at the beginning of transcription. The fragments produced with single-digoxigenin labeled primers generated torsionally unconstrained tethers for the following transcription assays.

### 4.3 Microchamber Preparation and Assembly of Transcription Tethers

Microchambers were assembled with laser-cut parafilm gaskets sandwiched between two glass coverslips [41]. The volume of a microchamber was about 10 µL. Antidigoxigenin (Roche Diagnostics) was introduced to coat the inner surface of the chamber at a concentration of 8 µg/mL in PBS for 90 minutes at room temperature. The surface was then passivated with Blocking buffer (PBS with 1% caesin, Gene-Tex) for 20 minutes at room temperature. Transcription tethers were assembled by mixing 30 nM of biotinylated RNAP holoenzyme and 3 nM linear DNA template in Transcription Buffer (20 mM Tris glutamate pH = 8, 50 mM potassium glutamate, 10 mM magnesium glutamate, 1 mM DTT, 0.2 mg/ml casein) and incubated 20 min at 37°C. Afterward, 50 µM ATP, UTP, GTP (NewEngland Biolabs), and 100 µM GpA dinucleotides (TriLink) were added to the solution and incubated for additional 10 min at 37°C to allow the ternary complex to initiat transcription and stall at the first G in the template. The solution of ternary complex was diluted to a final concentration of 250 pM RNAP:DNA complex, flushed into the passivated microchambers, and incubated for 10 minutes. Then, 20 µL of streptavidin-coated superparamagnetic beads (diluted 1:100 in Transcription Buffer; MyOneT1 Dynabeads, Invitrogen/Life Technologies) were flushed into microchambers to attach beads to biotinylated RNAP stalled on the DNA. Excess superparamagnetic beads in solution were then flushed out with 50 µL Transcription Buffer.

### 4.4 Magnetic Tweezers Assays

Magnetic Tweezers were used to observe the dynamics of transcribing ECs by recording the real-time changes in bead height. MTs consist of a pair of permanent magnets positioned above the microchamber that can be translated along the optical axis of a microscope to vary the strength of the magnetic field. The magnitude of tension can be calibrated from the lateral Brownian motion of the bead and the length of the DNA tether. Roadblock proteins were diluted to 20 nM (LacI) and 45 nM (EcoRI Q111) for optimal binding. The concentrations of LacI proteins were chosen based on previous looping experiments [42]. The concentration of EcoRI Q111 was selected by titrating the binding efficiency at different protein:DNA ratios from AFM images of EcoRI Q111:DNA complexes. The selected EcoRI Q111 concentration resulted competent on-site bindings with minimal off-site bindings in AFM images (data not shown). The diluted roadblock proteins were flushed into microchambers and incubated for 10 minutes at room temperature prior to recording. Excess roadblock proteins were then flushed out of the chamber with 40 µL of Transcription Buffer to limit binding to a single roadblock protein. After 3 minutes of recording, a complete set of NTPs was added into the chambers to resume transcription of CTP-starved ECs. Recordings lasted at least 30 minutes for LacI-O1/O2 roadblocks, and more than one hour for the observation of indefinitely stalled ECs.

Since high salt concentrations prevent the assembly and promoter escape of transcription complexes, Transcription Buffer with 50 mM [KGlu] was used to assemble halted transcription complexes. Then, for the high salt experiments, NTPs were flushed into microchambers in high salt Transcription Buffer (500 mM [KGlu]), mixing with normal salt Transcription Buffer in which halted ECs had been prepared, to give an overall concentration *∼*150 mM [KGlu].

### 4.5 Data Analysis

Prior to analyzing the data, we inspected the recordings to remove intervals with unusual signal fluctuations which may result from buffer addition or temporary tracking failures. Data in these intervals, mostly during the introduction of NTPs or GreA, were replaced with NaN values and discarded from further analysis. Next, we applied a Bayesian step change detection algorithm to the noisy raw data and extracted step-wise changes in tether lengths. The method minimizes the cost function

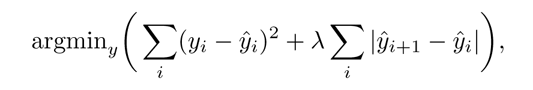

where *y*^ is the smoothed time series and *y* is raw time series. The method, which is described in detail elsewhere [43], reliably produced monotonic traces. The smoothed data consists of flat intervals separated by sharp jumps, and is well suited for detecting pauses. In theory, the positions of transcribing ECs along DNA templates can be calculated from the end-to-end distance of the tether knowing the force on the tether and the worm-like chain parameters of the DNA template. However, since there are small uncertainties in the force magnitude from tether to tether due to variation in bead size and magnetic susceptibilities, each individual trace was linearly scaled to align the prominent dwell times at promoter, roadblock and terminator positions. For this purpose, we converted the smoothed, monotonically increasing or decreasing traces to dwell time histograms. Since the smoothed traces change in a strictly step-wise manner, the histograms consist of sharp peaks representing the duration at pause sites. Next, we used expansion *a* and shift *b* factors to rescale the histograms, generating a new set of histograms *S^′^* = *a ∗ S* + *b*, to produce the histogram with maximum pause durations at the promoter, roadblock and terminator positions. Since the start point of transcription was set to 0 and always presents a significant pause, the shift factor was effectively zero. The best-fit values of expansion factor *a* ranged from 0.75 to 0.95, depending on the force magnitude in the experiments. Pauses within ±20 nm of the roadblock sites in the rescaled histograms were treated as roadblock-induced pauses.

### 4.6 Model Analysis

We used the Monte Carlo method to simulate pause times at roadblocks. The algorithm is shown in Algorithm 1.

Assuming that, under assisting force, RNAP passively waits for spontaneous dissociation of the roadblock, we postulated the values of *k*_3_ to be approximately equal to the inverse of the pause lifetimes of RNAP in front of the roadblock. The effects of assisting force and addition of GreA were included by setting the backtracking rate *k*_1_ = 0 and the backtracking recovery rate *k*_2_ = inf, respectively.

The roadblock dissociation rate *k*_3_ was determined by the pause lifetimes under assisting force experiments. The backtracking rate *k*_1_, recovery rate *k*_2_ and the probability of dislodging a roadblock, *P* 1, were fitted from the experimental pause lifetime distribution, with the assumption that *k*_2_ = 0.46 *k*_1_, which is derived from the energy profile of the transcription complex at the roadblock site.

## Supplementary information

Supplementary Figures S1-S4 available.

## Acknowledgments

LacI was a generous gift from Kathleen Matthews, Rice University. Plasmids for these experiments were created by Derrica McCalla and Zuzsanna Vörös. This work was supported by the National Institutes of Health (NIH) grants R01 GM084070 and R35GM149296 to LF and R01 GM067153 to IA.

## Author Contributions

J. Q. developed the experiments, collected and analyzed data, validated the model, and wrote the manuscript. A. C. helped collecting EcoRI Q111 roadblock data. W. X. and Y. Y. performed initial experiments and collected the LacI-O1 data in absence of tension. B. W. prepared proteins. I. A. supervised the preparation of proteins, performed co-immunoseparation analysis of the LacI-RNAP interaction, and participated in writing of the manuscript. D. D. designed the plasmids and with L. F. led the project, participated in the writing of the manuscript.

## Competing Interests

The authors declare no competing interests.

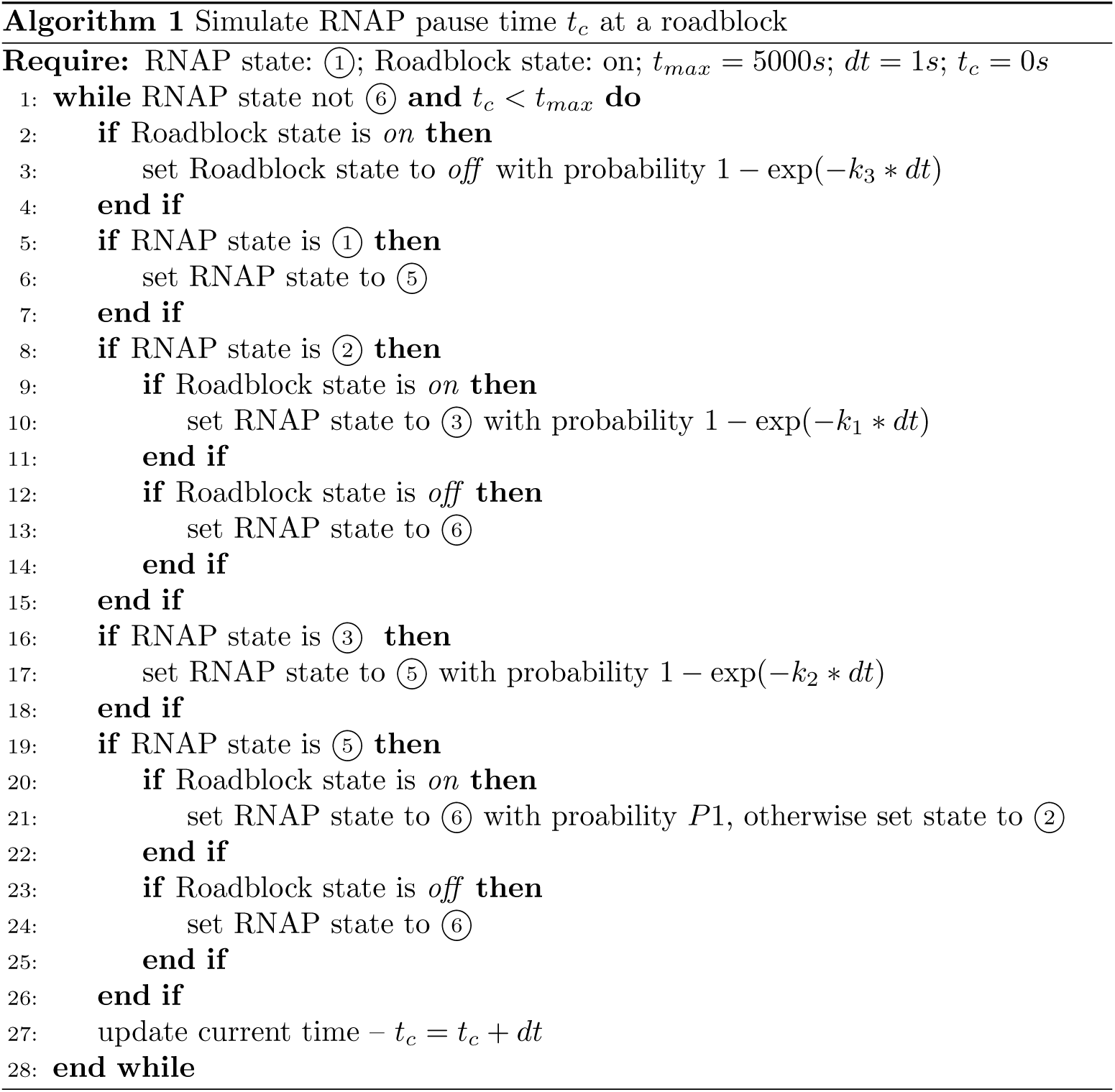

## Supplementary Information

**Fig. S1.**
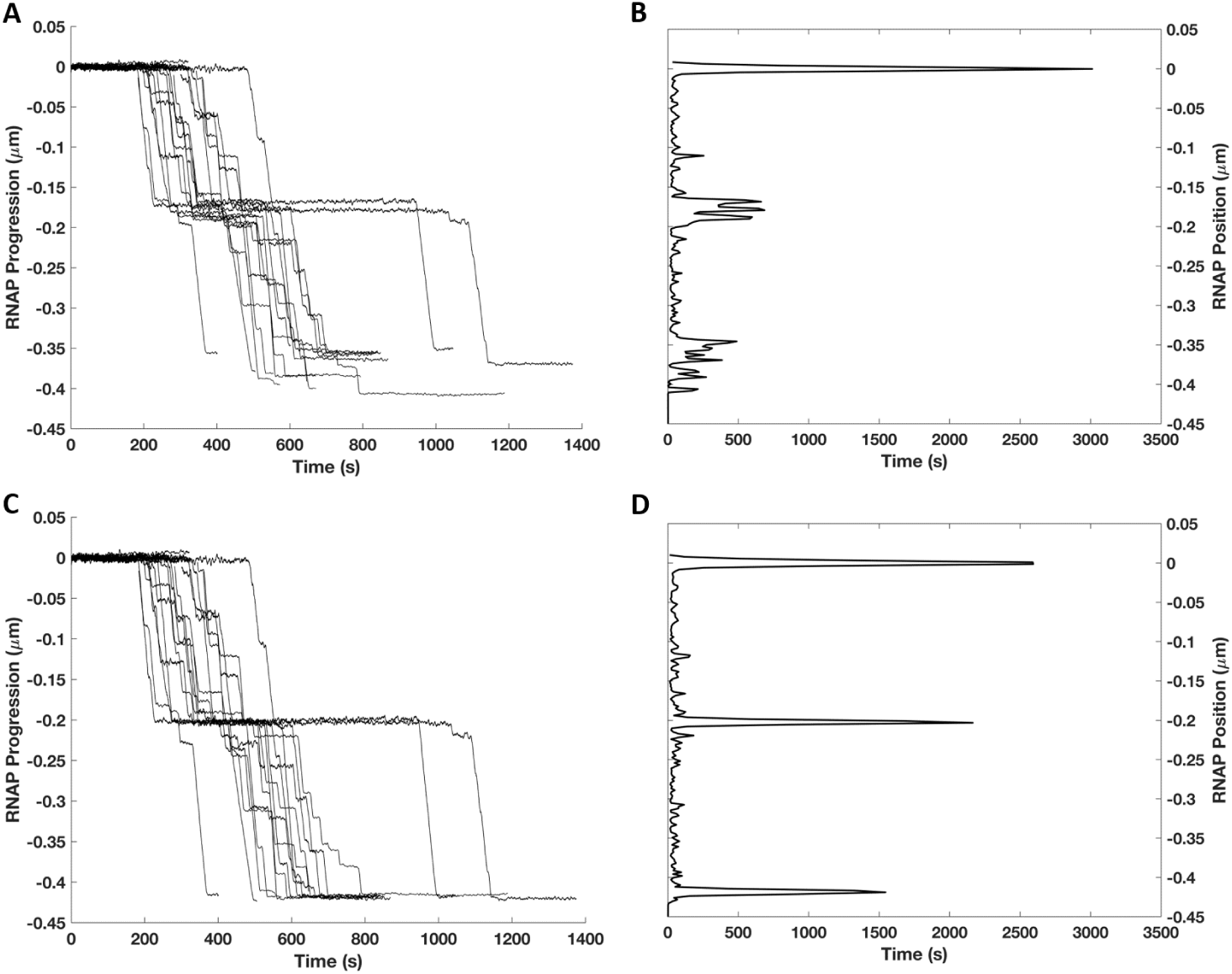
Processing of real-time transcription template length data. (**A**) A collection of transcription records under -2 pN (opposing) force. (**B**) The stacked histogram of the dwell times in the raw transcription records exhibits pauses dispersed around the roadblock and terminator positions before rescaling. (**C**) The same collection of transcription records in (**A**) are shown after rescaling to remove the effects of force variations that alter extension. (**D**) The dwell time histogram of the processed transcription records in (**C**) clearly shows peaks at promoter, operator and terminator sites.

**Fig. S2.**
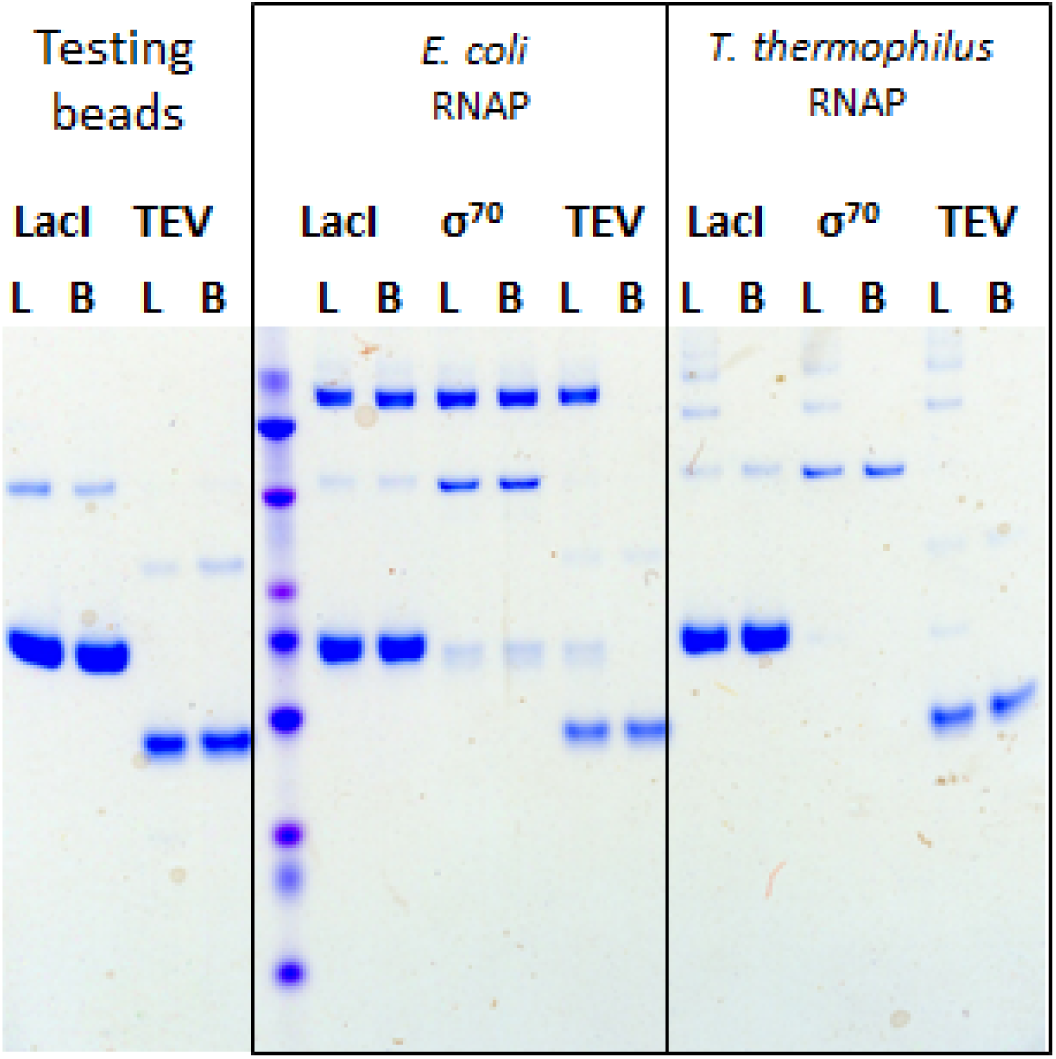
Co-immunoseparation of LacI with RNAP. (left) His-tagged LacI or TEV protease efficiently partitioned with Talon Dynabeads; L, loaded; B, bound. (middle) Untagged RNAP incubated with His-tagged LacI or *σ*70, but not with TEV, efficiently co-partitioned with Talon Dynabeads. (right) T. thermophilus RNAP incubated with LacI, *σ*70, or TEV did not co-partition with Talon Dynabeads.

**Fig. S3.**
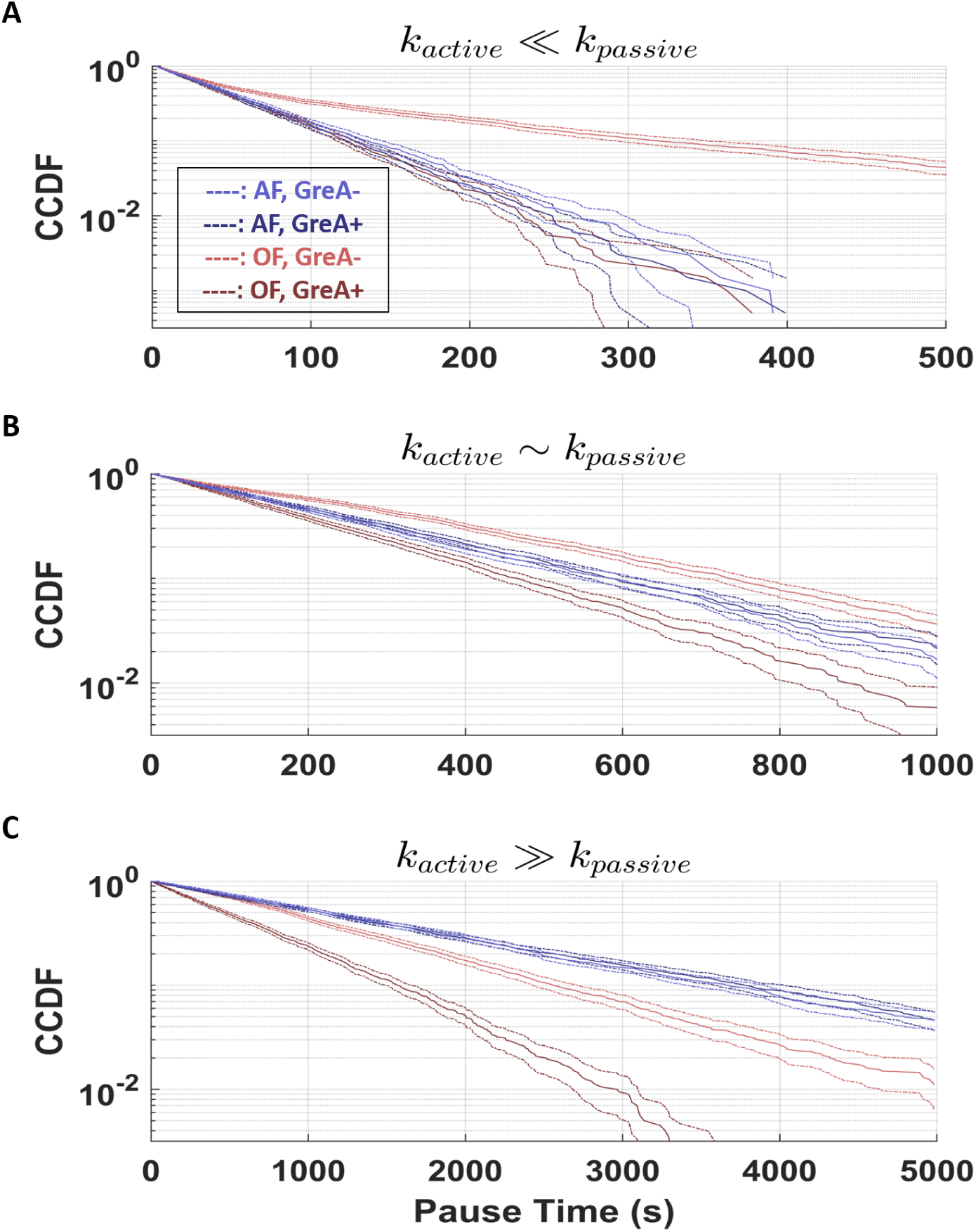
Simulation results. (**A**) In the regime *k*_active_ *≪ k*_passive_, the model produced complementary cumulative distribution functions (CCDF) showing *τ*_OF,GreA-_ *> τ*_OF,GreA+_ *∼ τ*_AF,GreA-_ *∼ τ*_AF,GreA+_. (**B**).In the regime *k*_active_ *∼ k*_passive_, the model produced CCDF showing *τ*_OF,GreA-_ *> τ*_AF,GreA-_ *∼ τ*_AF,GreA+_ *> τ*_OF,GreA+_. (**C**). In the regime *k*_active_ *≫ k*_passive_, the model produced CCDF showing *τ*AF,GreA-*∼ τ*AF,GreA+ *> τ*OF,GreA-*> τ*OF,GreA+.

**Fig. S4.**
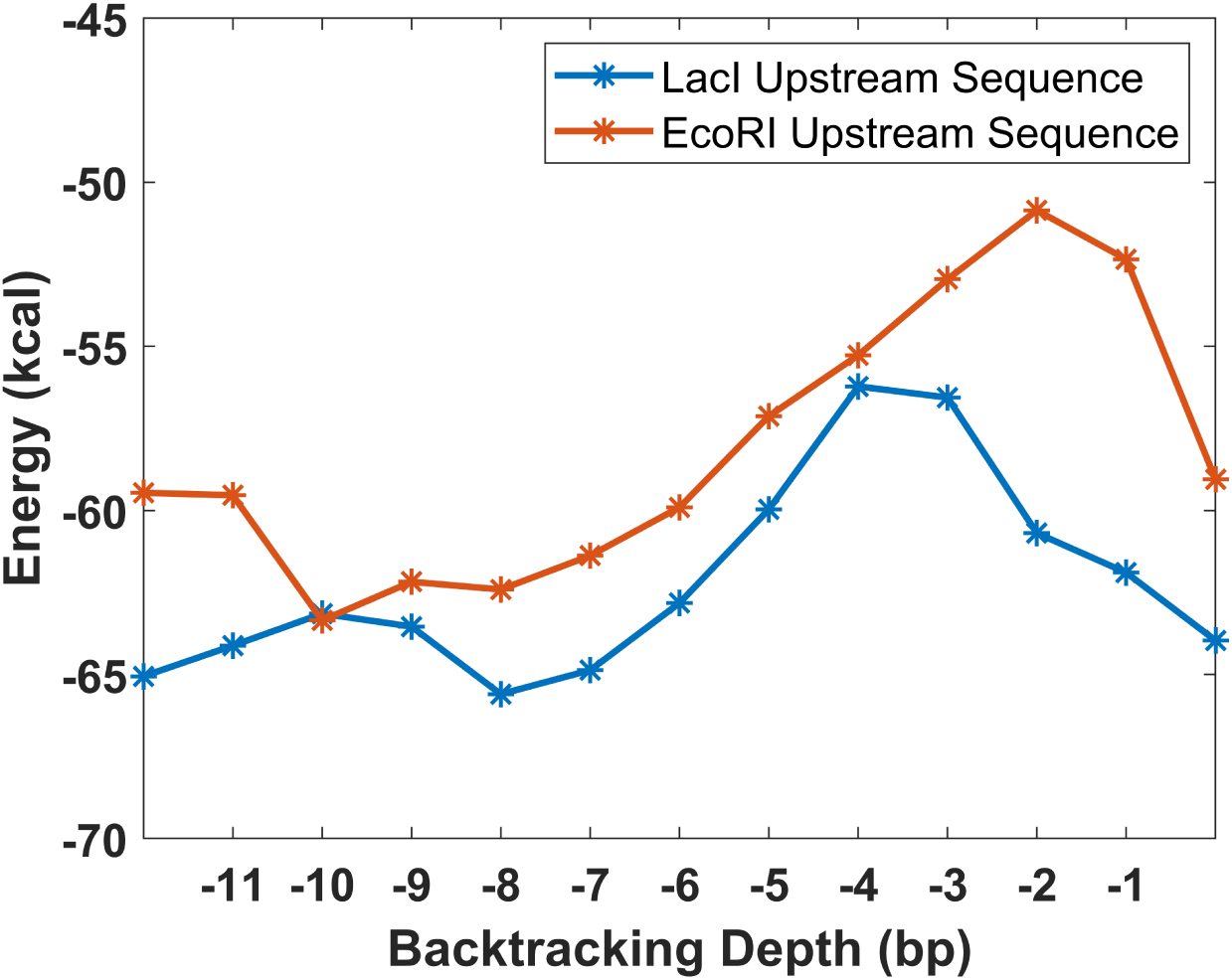
The free energy of ECs was calculated during backtracking upstream of LacI and EcoRI roadblocks as previously described [35].

